# Beyond infinite sites: Generalized ABBA-BABA statistic for deeper phylogenies

**DOI:** 10.64898/2026.07.06.736715

**Authors:** Chao Zhang, Rasmus Nielsen

## Abstract

The Patterson’s *D* statistic detects gene flow from ABBA-BABA site patterns, but its biallelic site patterns fail under deeper divergences where multiple hits cause false positives. We propose two extensions, *D*^+^ and *D*^*^. Both incorporate multiallelic site patterns to reduce saturation bias under JC and F84 model. Simulations show that *D*^+^ and *D*^*^ both remain correctly null under all conditions and detect gene flow effectively, with distinct advantages: *D*^+^ guarantees non-negativity of the denominator, while *D*^*^ provides greater robustness when mutation rates vary across genomic regions. The source code and binary files are publicly available at https://github.com/chaoszhang/ASTER.

## Introduction

Gene flow and hybridization are critical factors in evolutionary processes. The ABBA-BABA statistic, also known as Patterson’s D statistic (5; 3; 11), is a widely utilized method for detecting gene flow and hybridization in population genetic studies (5; 13; 17; 8; 2; 7; 14), and has been further explored by many theoreticians (10; 15; 12; 16; 6; 9). The D statistic operates by counting specific patterns of biallelic sites within aligned genomes of four taxa (a quartet). For a quartet phylogeny (((*T*_1_, *T*_2_), *T*_3_), *T*_4_), the counter *c*_*ABBA*_ represents the number of biallelic sites where taxon *T*_1_ matches taxon *T*_4_, and taxon *T*_2_ matches taxon *T*_3_. Conversely, the counter *c*_*BABA*_ represents the number of biallelic sites where taxon *T*_1_ matches taxon *T*_3_, and taxon *T*_2_ matches taxon *T*_4_. The D statistic is defined as:

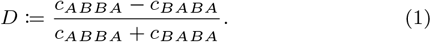

Under the null hypothesis of no gene flow, where genome-wide discordance is solely due to incomplete lineage sorting (ILS) within the multi-species coalescent (MSC) model and assuming the infinite-sites model (ISM), the expected values of *c*_*ABBA*_ and *c*_*BABA*_ are equal, resulting in *D* = 0. Therefore, the D statistic can be used to detect potential gene flow and hybridization by identifying quartet phylogenies that significantly deviate from this null hypothesis. However, the D statistic may not reliably detect gene flow and hybridization between distantly related taxa because the assumption of the ISM does not hold in deeper phylogenies. Our simulations further indicate that the D statistic can produce misleading results under such conditions.

In this article, we (i) present generalizations of the D statistic (referred to as D+ and D*), which extend its null hypothesis to the Jukes-Cantor (JC) model and the F84 model (4) of sequence evolution; (ii) prove that *D*^+^ = 0 and *D*^*^ = 0 under the null hypothesis of no gene flow and ILS within the MSC model, regardless of the depth of the phylogeny; and (iii) demonstrate through simulations that D+ and D* outperform the traditional D statistic in detecting gene flow and introgression in deeper phylogenies.

### Generalized D statistic

In this section, we define two ways of naturally extending D statistic (namely D+ and D* statistic), to JC model and F84 model, by including counters for multiallelic sites and applying proper weighting schemes. The D+ statistic, where

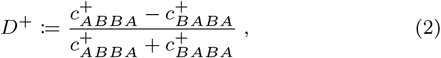

preserves the guaranteed non-negativity of its denominator but may be less effective for locating introgressed loci when mutation rate varies significantly among loci. The D* statistic, where

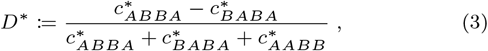

is robust to mutation rate variation among loci but at a risk of a negative denominator. We define 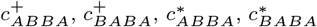 and 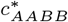 for JC and F84 model below.

#### Remark 1

As 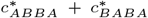 can easily become negative especially when ILS is very low, the denominator of (3) is defined as 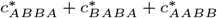 instead (Table 1).

**Table 1.**
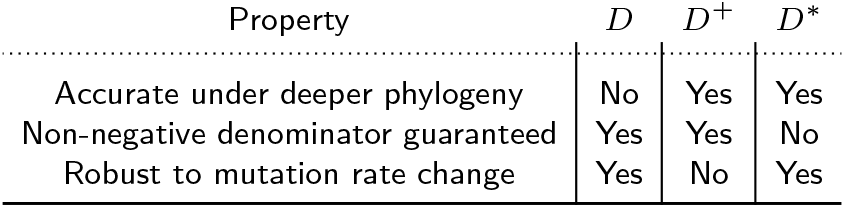
Statistics compared.

### Jukes-Cantor model

Additional counters in Figure 1 are introduced for each quartet phylogeny (((*T*_1_, *T*_2_), *T*_3_), *T*_4_). For example, *c*_*ABCA*_ denotes the number of sites where species *T*_1_, *T*_2_, and *T*_3_ have distinct nucleotides, and species *T*_1_ and *T*_4_ share the same nucleotide.

**Figure 1.**
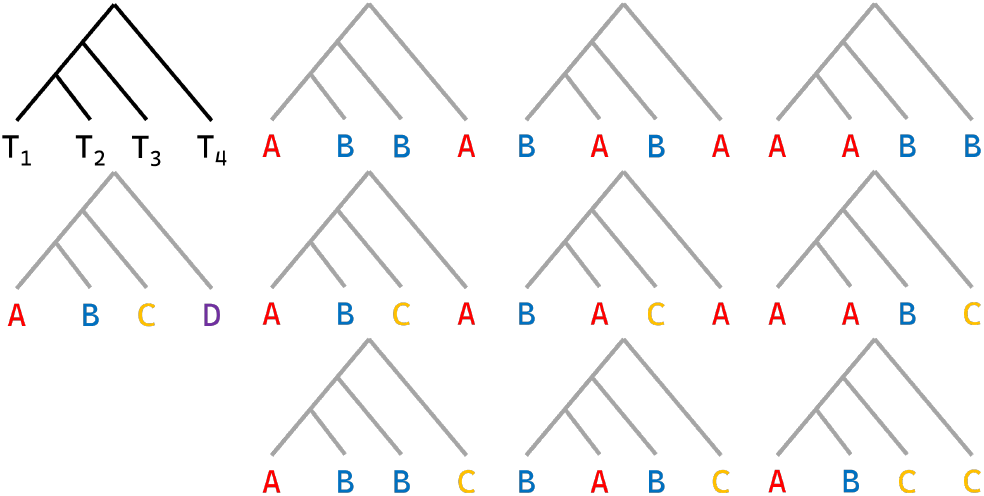
The extended set of site pattern counters for Jukes-Cantor model. Species trees represent site patterns in counters *c*_*ABBA*_, *c*_*BABA*_, *c*_*AABB*_, *c*_*ABCD*_, …, and *c*_*ABCC*_ . Different letters represent distinct nucleotides in a site pattern.

#### Definition 1

Counters for D+ statistic under JC model are defined as

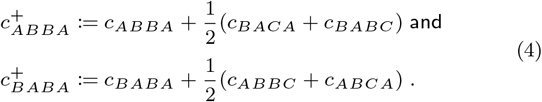

#### Remark 2

By the definition above, 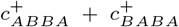 the denominator of (2) is non-negative. For the same reason, 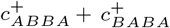 is non-negative for F84 model.

#### Theorem 1

Under the null hypothesis of quartet phylogeny under MSC+JC model, counters in Definition 1 satisfy that

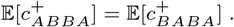

#### Definition 2

Counters for D* statistic under JC model are defined as

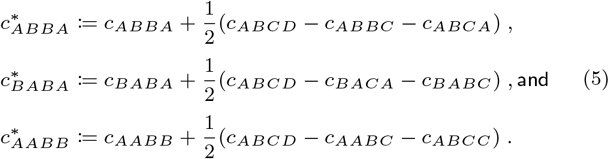

#### Remark 3

Under ISM, counters 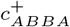 and 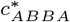 both reduce to *c*_*ABBA*_; counters 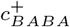 and 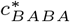 both reduce to *c*_*BABA*_.

#### Theorem 2

Under the null hypothesis of quartet phylogeny under MSC+JC model, counters in Definition 2 satisfy that

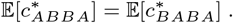

#### Theorem 3

Under JC model and assuming no ILS, counters in Definition 2 satisfy that

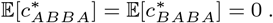

### F84 model

The definitions for D+ and D* statistic under F84 model are similar to ones under JC model, but counters 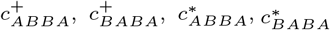 and 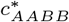 rely on more complicated patterns and weights depending on equilibrium frequencies to differentiate transitions versus transversions and weak versus strong hydrogen bonds (see Fig. S1). In practice, the equilibrium frequencies are estimated from the data.

#### Theorem 4

Under the null hypothesis of quartet phylogeny under MSC+F84 model, counters 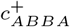and 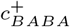 in Figure S1 satisfy

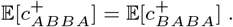

#### Theorem 5

Under the null hypothesis of quartet phylogeny under MSC+F84 model, counters 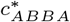 and 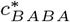 in Figure S1 satisfy that

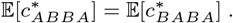

#### Theorem 6

Under F84 model and assuming no ILS, counters 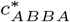 and 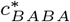 in Figure S1 satisfy that

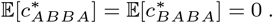

### Benchmarking via simulation

We simulated 4,800 replicates across four topologies, two mutation models, three divergence depths, and two mutation rates under the MSC model using `msprime` (see Supplementary Methods for the full factorial design). Tables S1–S3 in the Appendix present the full *Z*-score matrices and raw statistics for all conditions. We focus here on the key patterns.

### Null distribution under no gene flow

#### Classical *D* inflates with depth

Under the unbalanced tree (*utree*) with JC69, the *Z*-score of *D* grows from 0.0 *±* 1.2 at *t* = 1 to 5.1 *±* 1.7 at *t* = 5 (*µ* = 0.005) and reaches 79.7 *±* 19.0 at *t* = 25. The balanced tree (*btree*) shows the same trend: *D* reaches 87.4 *±* 24.1 (JC69, *t* = 25, *µ* = 0.005) and 90.9 *±* 29.1 (GTR, same depth and rate). At shallow depths (*t* = 1), *D* is correctly centered at zero across all models (|*Z*| *<* 0.5). The inflation is substantially worse at the higher mutation rate, confirming that multiple hits—not tree depth per se—drive the false-positive signal.

#### *D*^+^ **and** *D*^*^ **remain correctly null**

In contrast, *D*^+^ and *D*^*^ remain centered at zero across all null conditions. For *utree* under JC69 at *t* = 25, *µ* = 0.005, *D*^+^(F84) = −0.1 *±* 1.2 and *D*^*^(F84) = −0.1 *±* 1.2; under GTR the corresponding values are −0.5 *±* 1.1 and −0.5 *±* 1.1. Across all 24 null conditions (2 topologies *×* 2 models *×* 3 depths *×* 2 rates), |*Z*| never exceeds 0.5 for *D*^+^(F84) or *D*^*^(F84).

#### F84 is necessary

Under GTR, both *D*^+^ and *D*^*^ computed with the JC69 counting rule produce false positives at deep divergences, because unequal base frequencies violate the assumption of equal nucleotide composition. For *utree* GTR at *t* = 25, *µ* = 0.005: *D*^+^(JC) = 30.7 *±* 8.5 and *D*^*^(JC) = 30.8 *±* 8.5, indistinguishable from a genuine gene-flow signal. The F84-weighted versions correct this entirely: *D*^+^(F84) = −0.5 *±* 1.1 and *D*^*^(F84) = −0.5 *±* 1.1. This demonstrates that unequal base frequencies require explicit modeling: the JC69 counting rule, which assumes equal nucleotide frequencies, produces false positives under GTR, whereas the F84-weighted versions correct the bias (Fig. 2).

**Figure 2.**
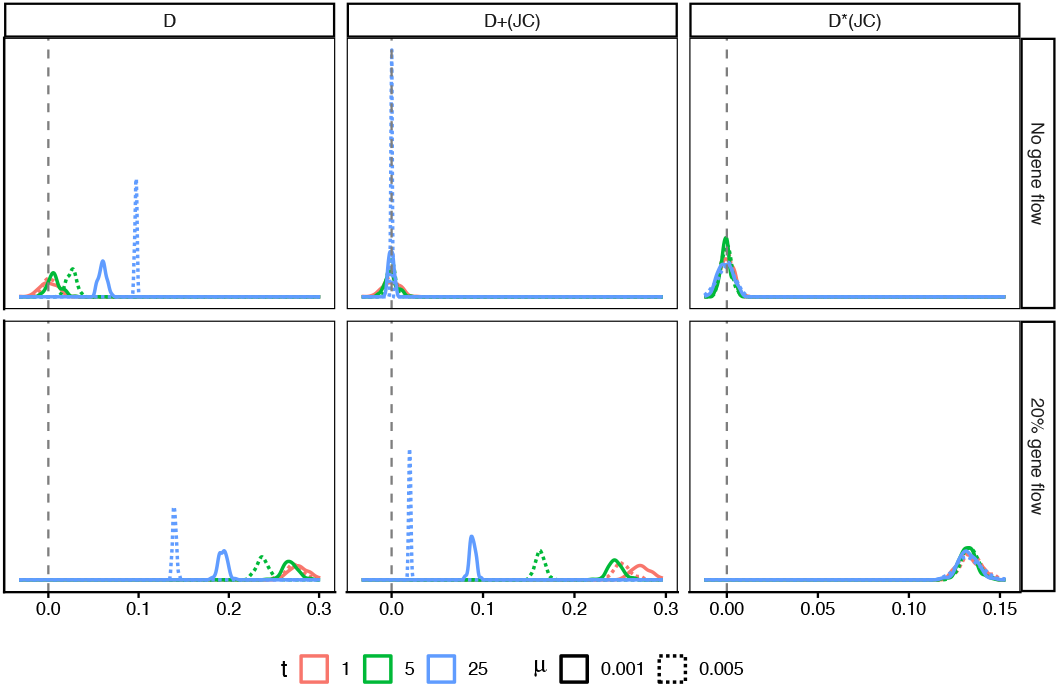
Distributions of *D, D*^+^(*JC*), and *D*^*^(*JC*) under JC69 model, for unbalanced tree (utree, no gene flow) and unbalanced network (unet, 20% gene flow). Solid lines: *µ* = 0.001; dotted lines: *µ* = 0.005. Columns correspond to divergence depths *t* = 1, 5, and 25. At *t* = 25 (blue), classical *D* is offset from zero even without gene flow (false positive), whereas *D*^+^(*JC*) and *D*^*^(*JC*) remain centered at zero. Under gene flow, all three statistics detect the signal at *t* = 25. *D*^+^ and *D*^*^ both remain accurate under deeper phylogenies, each with a different trade-off: *D*^+^ preserves non-negativity of the denominator (Table 1) but shows mutation-rate dependence (solid vs. dotted lines differ), while *D*^*^ is nearly invariant to mutation rate at the cost of guaranteed non-negativity (see also Fig. S2).

### Detection of gene flow

Under the 20% introgression scenario, all statistics detect the signal (Tables S1, S2).

#### Separate mutation rates

When mutation-rate conditions are analyzed independently, *D*^+^ and *D*^*^ produce comparable *Z*-scores. For the unbalanced introgression network (*unet* ) at JC69, *t* = 25, *µ* = 0.001: *D*^+^(F84) = 22.2 *±* 7.5 and *D*^*^(F84) = 21.7 *±* 6.8; at *µ* = 0.005: *D*^+^(F84) = 21.0 *±* 5.7 and *D*^*^(F84) = 21.0 *±* 6.1. The balanced network (*bnet* ) produces similar results. Notably, *D* itself remains a powerful detector at fixed mutation rate (*Z* = 92.0 for *unet* JC69, *t* = 25, *µ* = 0.005), but its inflation under the null makes it unreliable in practice.

#### Pooled mutation rates reveal *D*^*^ advantage

When chromosomes simulated under *µ* = 0.001 and *µ* = 0.005 are pooled into a single 20-chromosome replicate—simulating the realistic scenario where mutation rates vary across genomic regions—a striking divergence emerges. Under *unet* GTR at *t* = 25, *D*^+^(F84) drops to *Z* = 6.6 *±* 0.3, while *D*^*^(F84) retains *Z* = 25.4 *±* 3.9, a 3.8-fold advantage. The effect is consistent across models and topologies (Tables S1 and S2): *D*^*^ retains roughly 2–4*×* the *Z*-score of *D*^+^ in pooled-rate conditions.

#### Raw *D*^*^ values are mutation-rate invariant

The raw (unstandardized) statistic values explain this difference. For *unet* GTR at *t* = 25, *D*^*^(F84) is essentially constant: 0.134 *±* 0.009 at *µ* = 0.001 and 0.133 *±* 0.008 at *µ* = 0.005. In contrast, *D* drops from 0.186 *±* 0.005 to 0.133 *±* 0.002, and *D*^+^(F84) drops from 0.127 *±* 0.007 to 0.029 *±* 0.002. Across all gene-flow conditions and mutation rates, *D*^*^ remains stable at 0.132–0.134, while *D* and *D*^+^ vary by factors of 2–5*×*. The *c*^*^ counters, which incorporate multiallelic site patterns under the F84 model, absorb rate variation: as saturation increases, ABBA^*^, BABA^*^, and AABB^*^ all decay proportionally, leaving the ratio unchanged—consistent with the prediction of Theorem 5 (Figs. 2 and S2).

## Conclusion and discussion

We have shown that classical *D* inflates under deep divergences due to the failure of the infinite-sites assumption, and that two simple extensions—*D*^+^ and *D*^*^—correct this by incorporating multiallelic site patterns with model-based weighting. Both statistics remain correctly centered at zero under the null across all conditions tested. *D*^+^ preserves the two-pattern denominator of *D*, guaranteeing non-negativity regardless of model choice, but its power drops when mutation rates vary across genomic regions. *D*^*^ absorbs mutation-rate variation, but the two-pattern sum 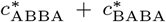 can become negative; adding 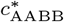 to the denominator reduces this risk substantially, though does not eliminate it. These complementary properties suggest that *D*^+^ is preferable when non-negativity is critical (e.g., per-locus scans), whereas *D*^*^ is better suited to genome-wide pooled analyses where mutation rates are heterogeneous. Future work should integrate *D*^+^ and *D*^*^ into Dsuite (9) to enable routine application, and extend the multiallelic-pattern framework to deeper statistics such as *F*_4_ and *F*_ST_, where saturation bias may be even more pronounced.

## Supplementary figures and tables

All supplementary figures and tables referenced in the main text are collected below.

**Figure S1.**
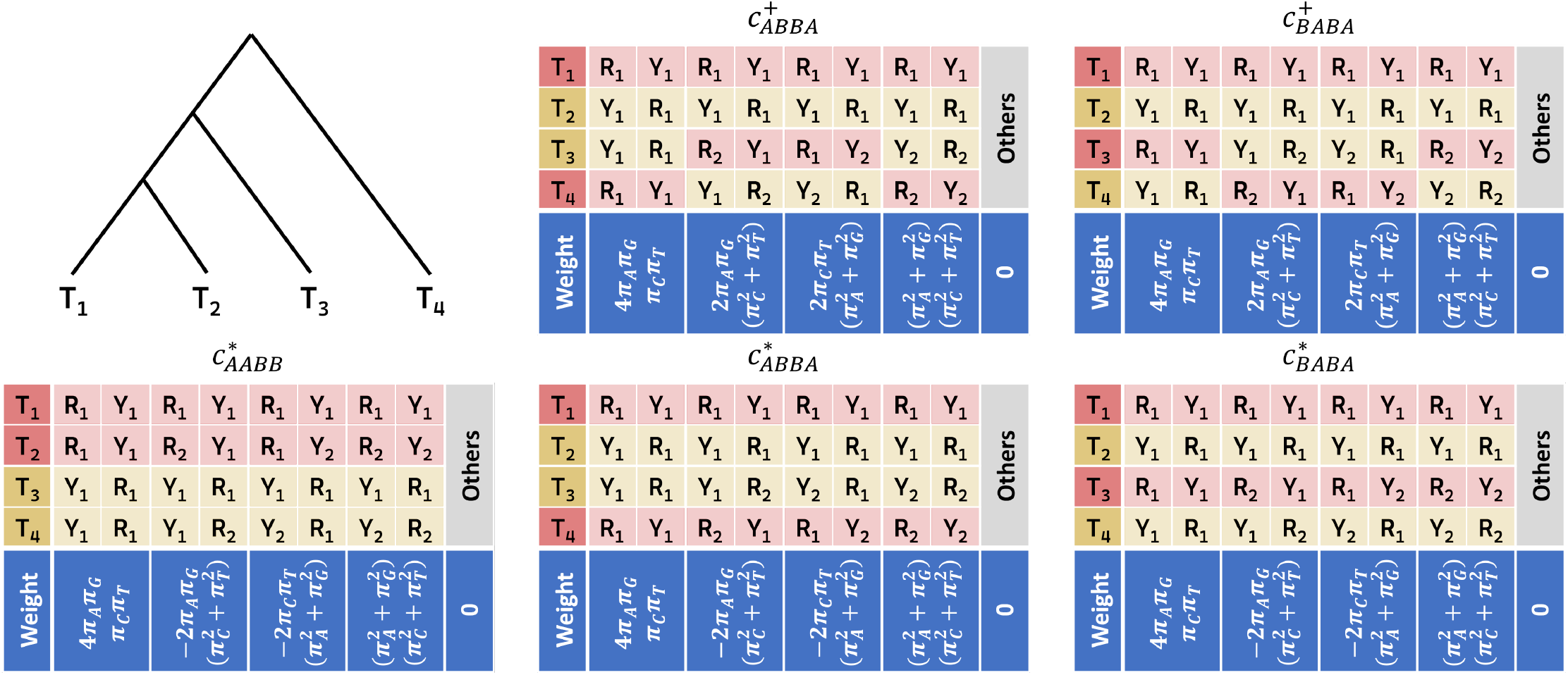
Site patterns for counters 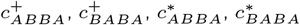, and 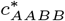 and their respective weights under F84 model. In site patterns, *R*_1_ and *R*_2_ denote different purines (A and G); *Y*_1_ and *Y*_2_ denote different pyrimidines (C and T). In weights, *π*_*A*_, *π*_*C*_, *π*_*G*_, and *π*_*T*_ denote equilibrium frequencies.

**Figure S2.**
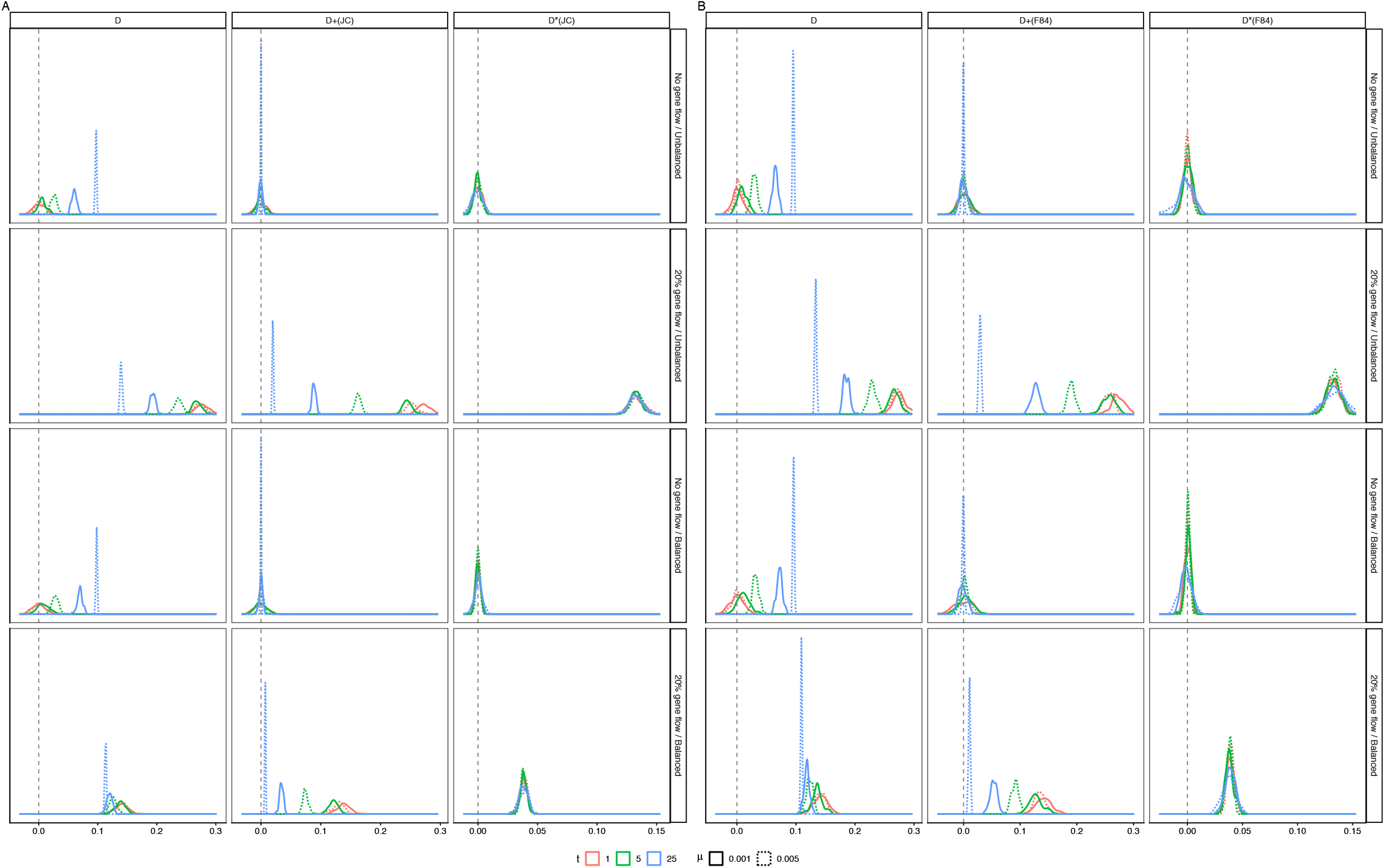
Distributions of *D, D*^+^, and *D*^*^ for all four topologies under JC69 (A) and GTR (B), columns correspond to the three statistics. Solid lines: *µ* = 0.001; dotted lines: *µ* = 0.005.

**Table S1.**
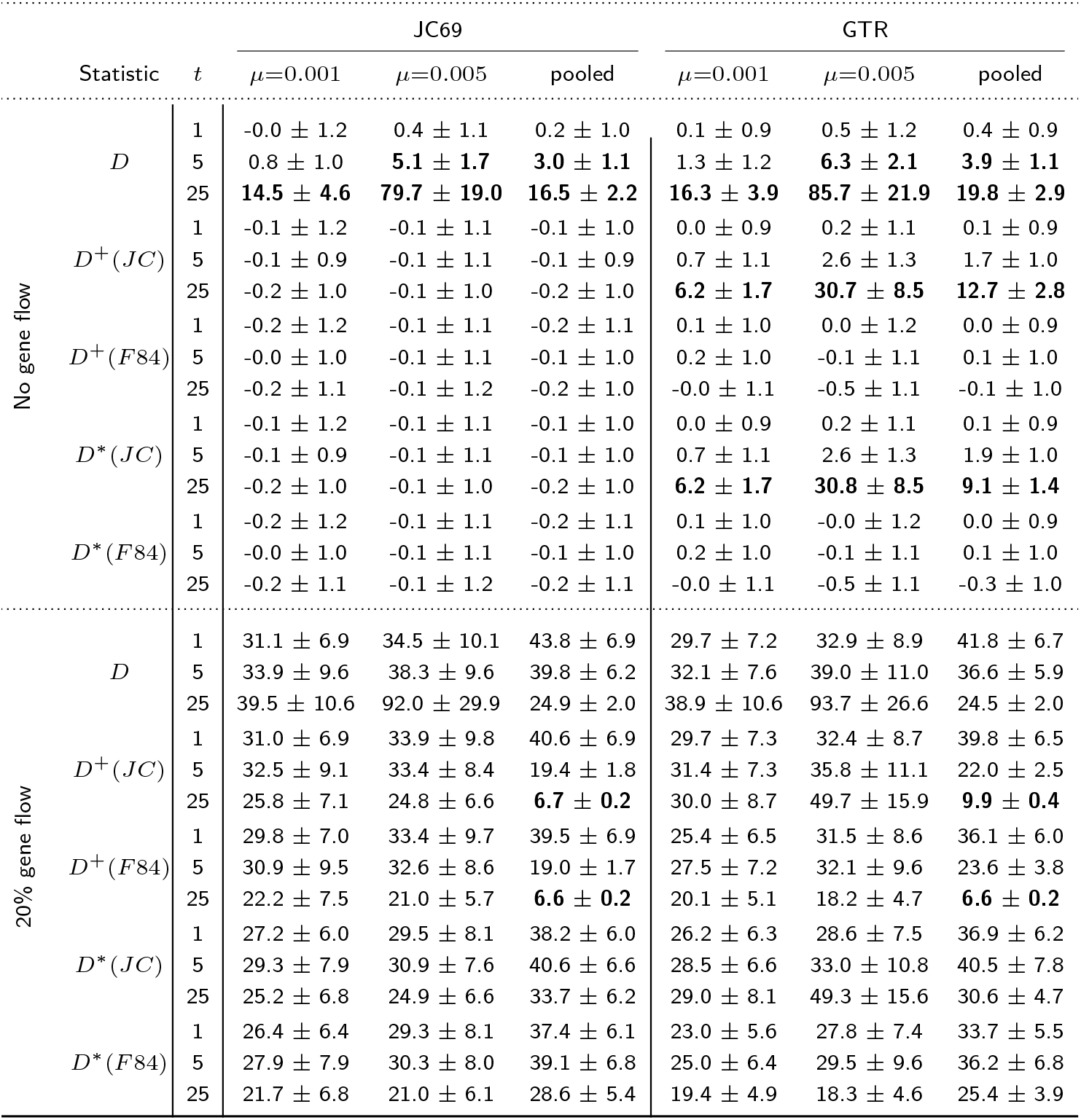
Mean and SD of per-replicate *Z*-scores for unbalanced topologies. Mean Z-values ≥ 3 in no gene flow cases and mean Z-values ≤ 10 in 20% gene flow cases are highlighted.

**Table S2.**
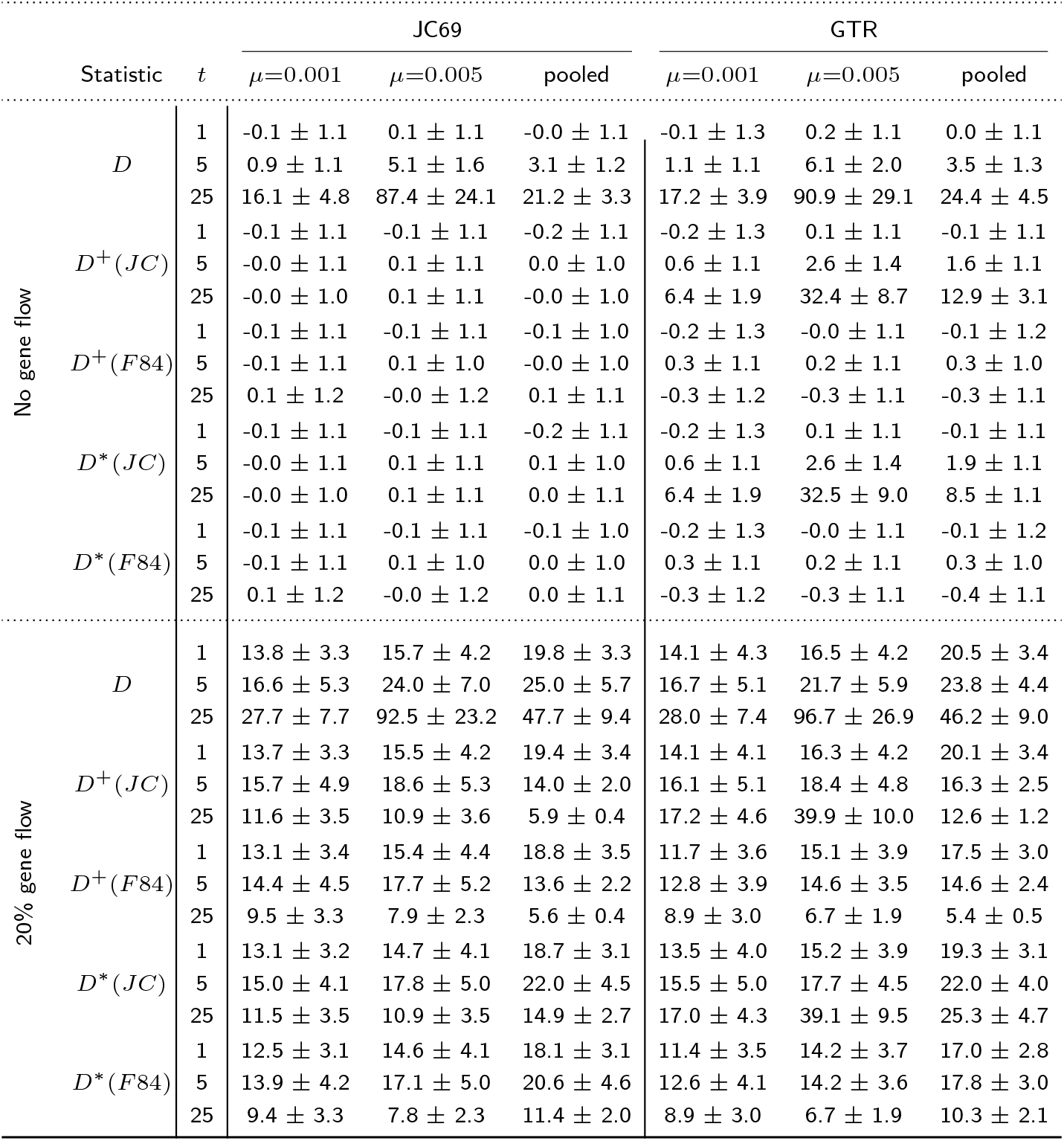
Mean and SD of per-replicate *Z*-scores for balanced topologies.

**Table S3.**
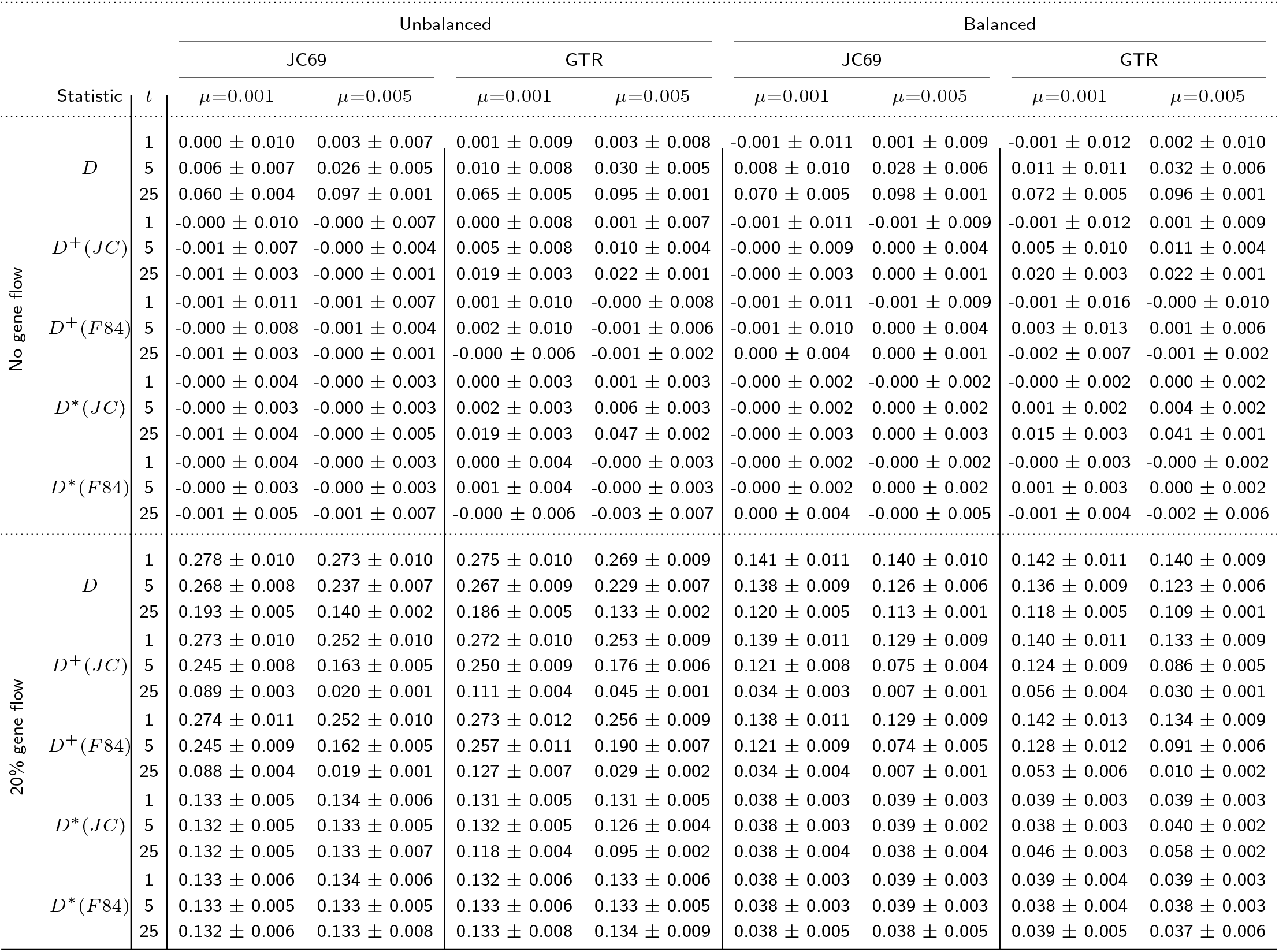
Mean and SD of per-replicate raw statistic values.

## Proofs

### Theorem 1

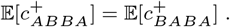

*Proof of Theorem 1*. We define

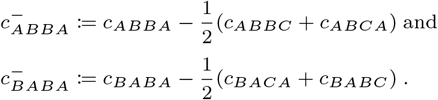

### Proposition 1

of (18) proved that under MSC+JC model,

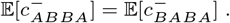

Since

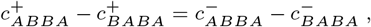

we conclude that under MSC+JC model,

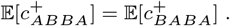

### Theorem 2

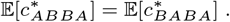

*Proof of Theorem 2*. Similarly, since

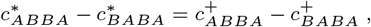

we conclude that under MSC+JC model,

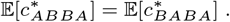

### Theorem 3

Under JC model and assuming no ILS, counters in Definition 2 satisfy that

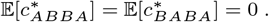

*Proof of Theorem 3*. For the species tree *S* of four leaves with topology *ab*|*cd*, let *l*_*a*_, *l*_*b*_, *l*_*c*_, *l*_*d*_ denote the terminal branch lengths, and *l*_*x*_ denote the internal branch length in substitution units. Under JC model:

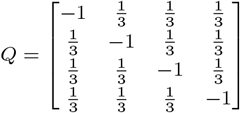

Let

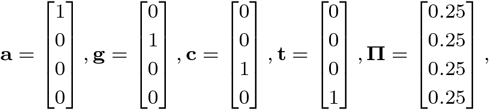

and

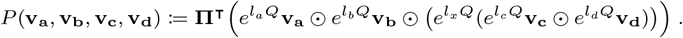

By definition,

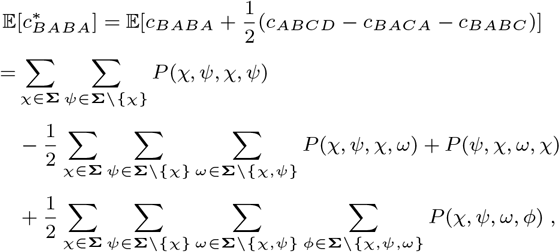

where **Σ** = *{***a, g, c, t***}*. By symmetry,

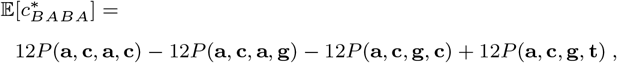

which is verified to be 0 using Mathematica (script provided in Supplementary Material).

### Theorem 4

Under the null hypothesis of quartet phylogeny under MSC+F84 model, counters 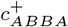 and 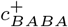 in Figure S1 satisfy that

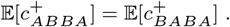

Proof of Theorem 4 is trivial after Proof of Theorem 5.

### Theorem 5

Under the null hypothesis of quartet phylogeny under MSC+F84 model, counters 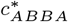 and 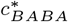 in Figure S1 satisfy that

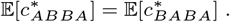

Proof of Theorem 5 can be found in Proposition 1 of (18) under MSC+F84 model.

### Theorem 6

Under F84 model and assuming no ILS, counters 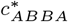 and 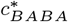 in Figure S1 satisfy that

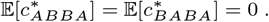

We omit the Proof of Theorem 6, as it follows the same proof of Theorem 3 and Proposition 2 of (18) under MSC+F84 model.

## Supplementary methods

### Simulation design

We evaluated *D, D*^+^, and *D*^*^ under a 4-taxon coalescent framework using msprime v1.4 (1). Each replicate simulated 10 independently evolving chromosomes of 10 Mbp under a recombination rate of 0.001 per bp per generation. We explored a factorial design:

- **Topology**: four ingroup phylogenies — unbalanced tree (*utree*), unbalanced network with 20% introgression from *T*_3_ to *T*_2_ (*unet* ), balanced tree (*btree*), and balanced network with 20% introgression (*bnet* ). All have the same outgroup *T*_4_ and identical divergence-order constraints.
- **Mutation model**: JC69 and GTR, with relative substitution rates *{*0.1, 0.3, 0.06, 0.12, 0.32, 0.1*}* and equilibrium frequencies 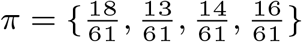
- **Divergence depth**: *t ∈ {*1, 5, 25*}*, measured in coalescent units from the root of the ingroup to the present.
- **Mutation rate**: *µ ∈ {*0.001, 0.005*}* per bp per 4*N*_*e*_ generations.
- **Replicates**: *N* = 100 per condition, for a total of 4 *×* 2 *×* 3 *×* 2 *×* 100 = 4,800 replicates and 1.92 TB of simulated sequences.

For each replicate, we pooled the 10 chromosome-level site-pattern counts before computing *D, D*^+^, and *D*^*^. For *D*^+^ and *D*^*^ under the F84 model, we used the JC69 formula (Definitions 1 and 2) as well as the F84-weighted formula (Fig. S1) to compare the JC69 counting rule with the full F84 base-composition correction.

### Statistical evaluation

For each condition and each statistic *S*, we computed the per-replicate *Z*-score 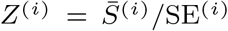, where 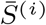 is the value of *S* pooled across 10 chromosomes of replicate *i*, and SE^(*i*)^ is the standard error of the chromosome-level values. We then report the mean and standard deviation of these *Z*-scores across the *N* = 100 replicates: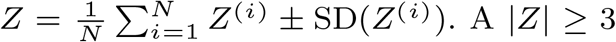 threshold flags significant deviation from the null expectation of zero. We additionally report raw statistic values as 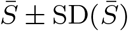 across replicates.

